# Herbivore pressure modulates soil multifunctionality in a Mediterranean landscape

**DOI:** 10.64898/2025.12.12.693896

**Authors:** Jorge F. Henriques, João Carvalho, Rita Tinoco Torres, Mariana Rossa, Joana M. Fernandes, Catarina Malheiro, Marija Prodana, Sara Peixoto, Eduardo Ferreira, Susana Loureiro, Jennifer Adams Krumins, Ramón Perea, Emmanuel Serrano, Rui G. Morgado

## Abstract

Ongoing global environmental changes demand cost-effective strategies to restore ecosystem functioning. Within this context, the reintroduction of large mammalian herbivores has emerged as a promising approach to recover ecological processes and foster self-sustaining, biodiverse ecosystems. However, the impacts of such initiatives remain poorly understood, particularly regarding their effects on soil functions and ecological processes. We tested whether large herbivore densities (wild horses (*Garrano* breed) and cattle (*Maronesa* breed, an ancient lineage close to aurochs) modulates soil multifunctionality (the simultaneous provision of multiple ecosystem functions) across habitats (grasslands, shrublands, and forests) and seasons (fall and spring) in a Mediterranean landscape. Using composite multivariate indices describing soil functions related to biogeochemical cycles (i.e., enzymatic activities) and physicochemical properties (pH, conductivity, organic matter, and nutrient loads), we found that soil enzymatic activities varied seasonally, being higher in spring, and interacted with habitat and herbivore pressure gradients. Habitat and herbivore pressure explained the variation in soil multifunctionality during spring, while in fall it was mainly driven by herbivore pressure. Enzymatic stoichiometry, particularly C:P ratios, strongly predicted soil fertility and multifunctionality, showing positive relationships in forest habitats and under herbivore pressure. A mechanistic approach confirmed that herbivore impacts on soil functioning operated primarily through changes in soil properties and nutrient cycling rather than direct effects. Mediterranean landscapes are rapidly changing, and our results highlight herbivore management as a key tool to sustain soil ecological processes under the rewilding framework.

**Highlights:**

- Large herbivores impacted soil functions and properties in different habitats
- Large herbivores significantly impacted soil nutrient cycling and multifunctionality.
- The magnitude of the effects was dependent on season and habitat
- Large herbivores indirectly impact soil ecosystem multifunctionality through changes in soil properties.

## Introduction

The accelerating pace of global changes has fostered profound discussions on strategies for effective landscape management. Among these, the reintroduction of large mammalian herbivores into areas from which they were extirpated has emerged as a cost-effective measure to restore ecological processes and promote the recovery of self-sustaining, and biodiverse ecosystems (Cromsigt et al. 2018; Schmitz et al. 2023). In theory, reintroduced herbivores may enhance biodiversity and ecosystem resilience (Bakker and Svenning 2018; Cromsigt et al. 2018) through cascading effects on biological communities (e.g., vegetation and soil). As ecosystem engineers, large herbivores can modulate ecological processes by shaping the structure and properties of terrestrial food webs above- and belowground (Ramirez et al. 2018). Yet, the ecological mechanisms underlying these effects are complex, reflecting the direct and indirect pathways within soil-plant-herbivore interactions (Castillo-Garcia et al. 2022). Large herbivores can directly shape the plant community structure aboveground through grazing, browsing, and trampling (Taboada et al. 2015; Sitters and Andriuzzi 2019), as well as the biogeochemical cycles belowground by changing nutrient loads through the excretion (e.g., urine and feces) and processing of plant material (Sitters and Andriuzzi 2019). These interactions are replicated across ecosystem compartments, giving rise to context-dependent and often indirect effects on soil functioning. Understanding these effects requires an integrative framework that simultaneously accounts for multiple soil functions.

The concept of soil ecosystem multifunctionality (EMF) provides such a framework, as it integrates several functions that underpin ecosystem services (Marshall et al. 2025). Nevertheless, this framework has not yet been sufficiently explored to evaluate the effects of large herbivores rewilding but its integrative value and the close relationship with the provision of multiple soil-based ecosystem services highlights this valuable approach (Svenning et al. 2024). While direct transposition to real-world policies is challenging (Garland et al. 2021), this framework has helped to unveil biodiversity-ecosystem functioning relationships, offered mechanistic insight into the ecological processes, and proven helpful in foreseeing impacts and proposing effective mitigation measures for soil conservation (McGranahan 2014). Whereas research on the impacts of large herbivores, including their impacts on soil, has been the subject of many studies in last three decades (Carvalho et al. 2025), significant gaps remain under the rewilding framework, particularly on integrating multivariate knowledge for establishing causal links to soil multifunctionality. The magnitude and direction of the effects of ungulates is contingent upon several factors, including species (namely, livestock vs wild ungulates; Marks et al. 2024), grazing intensity and duration (Abdalla et al. 2018), plant functional traits (Peco et al. 2012), climate, soil properties (Dodge et al. 2020; Lai and Kumar 2020), and vegetation types (McSherry and Ritchie 2013). As the impacts of ungulates are often expressed through indirect pathways, understanding the mechanisms driving soil responses is important for selecting appropriate indicators to assess perturbation effects on soil functions. In this sense, soil enzymes are key actors regulating organic matter decomposition and nutrient cycling (Raiesi and Salek-Gilani 2018), highlighting their relevance as indicators of soil multifunctionality (Garland et al. 2021). To disentangle the direct and indirect impacts of ungulates on soil multifunctionality, this study aims to understand how soil EMF varies due to changes in (i) ungulate herbivore pressure (absent, low, or high), (ii) habitat (grasslands, shrublands, and forests), and (iii) seasons (fall and spring), and whether the observed patterns are related to (iv) soil physical and chemical properties, or not.

We hypothesize that (H1) ungulates will impact soil multifunctionality through their effects on soil physical and chemical properties (Marks et al. 2024); (H2) these changes alter nutrient cycling (e.g., Barbero-Palacios et al. 2020)) and (H3) the magnitude of these impacts depends on habitat and season, with stronger effects on grasslands, during the dry season due to greater biomass removal. Overall, this study aims to provide a mechanistic understanding of how rewilding with ungulate herbivores shapes soil multifunctionality across contrasting ecological contexts.

## Material and Methods

### Study area

This study took place in the Faia Brava Reserve (FBR), a fenced protected area (40.9055°N,-7.0960°W; 857 ha) located in central Portugal, located within the Côa Valley (Figure 1). This is a typical Mediterranean landscape with steep slopes and rocky plains shaped by human activities such as agriculture and herding. It encompasses a range of Mediterranean-type habitats, including forests dominated by *Quercus rotundifolia* and *Quercus suber* (“montado”, 17.4%), thermo-Mediterranean scrubs (18.6%), European dry heaths (41.2%), pseudo-steppe with grasses and annuals (18.4%), and several natural and artificial temporary ponds. This system is classified as a hot-summer Mediterranean climate (Kottek et al. 2006), with temperatures ranging from na average of about 5°C in winter and 28°C in summer. Annual average precipitation is about 400-600 mm, with a pronounced summer drought. The FBR has been, since 2005, under a management regime to control shrub encroachment and promote landscape heterogeneity using herds of semi-wild large ungulates in an extensive regime. The herds include wild horses (Garranos breed) and cows (Maronesa breed), an ancient breed closely related to aurochs. These animals were equipped with GPS-GSM collars (DigitAnimal GPS Trackers) allowing monitoring of their movement and space use. Based on the location of individuals through time, the study area was classified accordingly (Figure 1).

**Figure 1.**
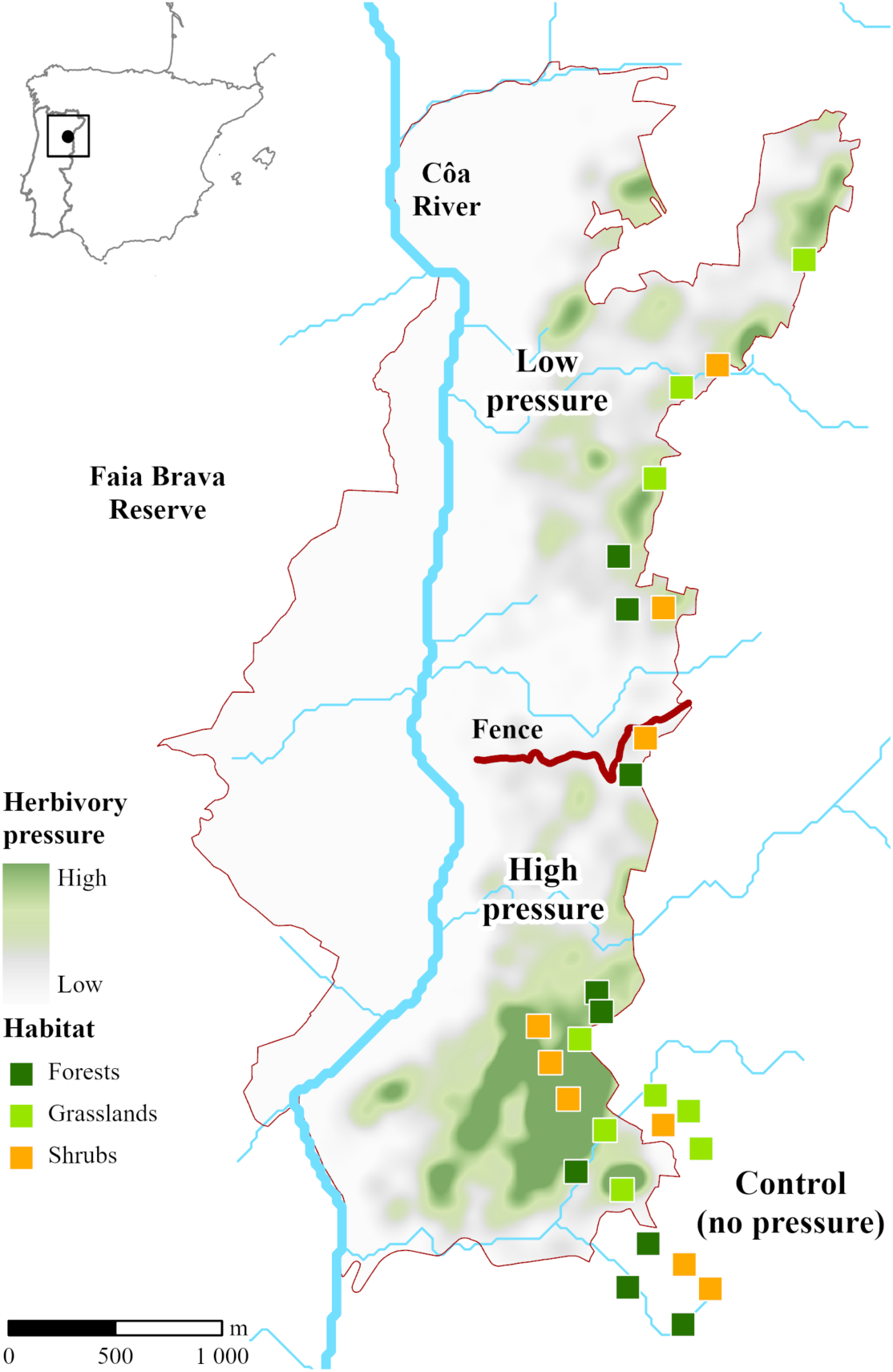
Ungulate herbivore distribution and pressure level in the study sites at Faia Brava reserve. Squares in light green represent grasslands, in yellow the shrublands and in dark green the forest plots.

### Experimental design

We established a full factorial experimental design with three types of habitats and three levels of ungulate herbivore pressure, each comprising three replicates, i.e., sites, (Figure 1). Two sampling plots were established at each site, each consisting of paired fenced and unfenced areas of 225 m² (15 x 15 m). This paired fenced/unfenced setup belongs to a long-term manipulation experiment initiated shortly before the first sampling campaign, with no significant differences observed between treatments at this early stage. In total, 45 sampling plots were sampled in two seasons (fall 2021 and spring 2022). During the implementation of this study, there were 35 wild horse and 35 cows, distributed throughout the reserve (1 animal p/ 7.5 hectares, i.e., 1 wild horse p/ 15 hectare and 1 cow per 15 hectare). Due to high aridity and the habitat characteristics of our study area, with high percentage of rocky outcrops, dense shrublands and sparse grasslands, these animal densities are above the carrying capacity of this system, and thus, the herbivore pressure is relatively high. Animal were equipped with GPS-GSM collars since 2019, which allowed us to assess herbivory pressure throughout the FBR and define sampling sites for our experimental treatments. Combining current occurrences and historical records, herbivore pressure treatments were designated as: (i) control (no grazing), (ii) low herbivore pressure (0.04 animal/ha) and (iii) high herbivore pressure (0.30 animals/ha) (Figure 1).

Regarding habitat types, the most spatially representative habitats within mediterranean landscapes were considered: i) grasslands, ii) thermo-Mediterranean shrublands (hereafter, shrublands) and forests of *Quercus* spp. (hereafter, forests). Candidate areas sharing the same topographic characteristics, such as slope, appearance, and roughness, were selected using a combination of satellite-aided surveys and field observation.

### Soil sampling and preparation for analysis

We collected soil samples in the 45 previously defined experimental plots using polyvinyl chloride (PVC) soil cores (0-7 cm depth). A total of 12 cores randomly distributed over the 225 m^2^ area were pooled for each sampling plot, ensuring that soil samples were representative of each sampled habitat. Soil samples were immediately placed in a refrigerated container and stored at 4 °C as soon as possible. Before analysis, moist soil samples were sieved to 2 mm to remove plant litter and roots. Between each sieving procedure, sieves were cleaned with a diluted sodium hypochlorite solution (3%) and thoroughly rinsed with water. After processing, each bulk was subsampled for physical and chemical properties, enzyme activities, and nutrients.

### Soil physical and chemical analysis

Soil pH and electrical conductivity (EC) were measured at a soil-water ratio of 1:5 according to standard procedures (ISO 1994). Soil moisture, i.e., gravimetric water content (GWC), was assessed by calculating the proportional weight loss of fresh soil (i.e., obtained directly from the field) after drying at 105 °C for 48 h. Organic matter (OM) content was determined by loss on ignition at 500 °C for 6 h. Weight loss due to organic matter atomization was estimated considering the oven-dried soil as the base (Schulte and Hopkins 1996). Bulk density (BD), i.e., the weight of a volume unit of soil, was assessed by weighing oven-dried soil in a container with a known volume (74 cm^3^). Total carbon (TC) and total nitrogen (TN) contents were measured by an elemental analyzer (VarioMax CNS; Elementar, Langelsbold, Germany). Total phosphorus (TP) was measured in *aqua regia*-digested soil samples with ICP-OES (Jobin Yvon Activa M), following standard procedures (ISO 11885). Soil organic carbon (SOC) was determined using the Walkley-Black colorimetric method (GLOSOLAN 2020). Hot water extractable fractions of carbon (HWEC) and nitrogen (HWEN) were extracted as described in (Ghani et al. 2003) and quantified using a TOC analyzer (Analytikjena multi N/C 100). Soil available phosphorus (BRAY-P) was determined using the Bray-1 colorimetric method following standard operating procedures (FAO 2021). Available cations (potassium - AK, magnesium - AMg, and Calcium - ACa) were extracted with ammonium acetate (NH₄CH₃CO₂) solution and quantified through Inductively Coupled-Plasma Atomic Emission Spectroscopy (ICP-AES, Horiba Jobin Yvon ULTIMA sequential ICP), according to the standard operating protocol (FAO 2022).

#### Soil enzymatic activities

Enzymatic activities were measured in soil samples collected from areas with different types of habitats and ungulate herbivory pressure levels to assess the contribution of microbial activity to nutrient cycling. The selected enzymes targeted key nutrient cycles: β-glucosidase (BG - carbon cycle), alkaline and acid phosphatase (AkP and AcP, respectively - phosphorus cycle), arylsulfatase (AS - sulfur cycle), urease (UR - nitrogen cycle), and dehydrogenase (DHA) as an indicator of overall microbial activity (Wolinska and Stepniewsk 2012). The activity of DHA was quantified by determining the amount of 1,3,5-triphenylformazan (TPF) released after incubating 3 g of fresh soil with a substrate solution consisting of 2,3,5-triphenyl tetrazolium chloride (3%) for 24 h at 37 °C (Tabatabai 1994). The activities of BG, AcP, AkP, and AS were measured as the amount of p-nitrophenol (PNP) released after incubating 0.5 g of fresh soil with the respective substrate solutions: i) p-nitrophenyl phosphate (0.05 M), ii) 4-nitrophenyl β-D-glucopyranoside (0.05 M) (Acros organics, 99% of purity), and iii) 4-nitrophenyl sulfate (0.05 M) (Acros organics, 99% of purity) (Dick et al. (1997). The incubations lasted for 1 h at 37 °C. AcP and AkP were measured using a modified universal buffer (MUB) at pH 6 and 11, respectively. UR activity was quantified as the amount of ammonium obtained after incubating 1 g of fresh soil with urea and borate buffer (pH 10) for 2 h at 37° C (Tabatabai 1994). The TPF, PNP, and urea concentrations were determined through photometric analysis at 485 nm, 410 nm, and 690 nm, respectively, using a microplate spectrophotometer (Thermo Scientific mod. Multiskan Spectrum). All results are expressed based on oven-dried (105 °C, 48 h) soil weight.

### Data analysis

#### Soil properties and multifunctionality indices

The statistical analyses and indices estimated within the scope of this work were conducted in R software (v4.4.1). Spearman rho correlation coefficient was used to test associations between soil functions, physical and chemical parameters, and other environmental variables (Figure S1). Normality was assessed through the Shapiro-Wilk normality test, and variables were transformed accordingly using a natural logarithm function (ln). To calculate the *soil ecosystem multifunctionality index,* a multivariate approach was used, based on the principal component analysis (PCA) scores of the standardized values (z-scores) of laboratory-measured soil functions (DHA, BG, AS, AcP, AkP, and UR) (Meyer et al. 2018). A soil *fertility index* was calculated similarly, but gathering a different set of measured soil indicators broadly known to be surrogates for plant primary productivity, e.g. (Moran et al. 2000; Amara et al. 2017; Munnaf and Mouazen 2021), including soil physicochemical properties, nutrients, and elements (TC, TN, TP, SOC, HWEC, HWEN, BRAY-P, AK, AMg, ACa, OM, GWC, pH, EC, BD). When conducted in parallel, these indices provide a holistic overview of soil quality, allowing benchmarking and validation (Gutierrez et al. 2024). Finally, the following indices were produced using targeted subsets of data: i) *carbon cycling index*, based on variables closely related to the carbon cycle (BG, HWEC, SOC, and TC); ii) *nitrogen cycling index*, with variables closely related to the nitrogen cycle (UR, HWEN, and TN); ii) *phosphorous cycling index*, with variables closely related to the phosphorous cycle (AcP, AkP, BRAY-P, and TP); and iv) *physicochemical properties index,* based on the soil properties capable of modulating nutrient availability and enzymatic activities (pH, EC, GWC, BD, AK, AMg, and ACa) (Table S1). Similarly, nutrient availability indices were considered as complementary approach (Appendix 1).

#### The effects of ungulate pressure and habitat

The effects of ungulate pressure, habitat, season, and their interactions on: i) soil enzymatic activities, ii) soil multifunctionality, and iii) other composite indices produced, were assessed using linear mixed effects model (LMMs), fitted using the lme4 package (Bates et al. 2007) in R software (v4.4.1). The predictors (fixed effects) were ungulate herbivory pressure [Control (no pressure), low and high pressure], season (fall and spring), and habitat (grasslands, shrublands, and forests). In all the models, the site identity was used as a random effect to account for background variation due to sampling locations, as there were two plots within each site. The relevance of the fixed effects was evaluated based on their inclusion in the best-fitting model, defined as the model with the lowest AIC value and differing by more than two units from any simpler model. (Burnham et al. 2011). Due to seasonal changes in soil properties, model selection was performed separately on each season’s datasets. Finally, to further explore the context-dependency associated with ungulate pressure and habitat, regression analysis and Pearson correlation coefficient was used to assess soil enzymatic and nutrients stoichiometry, as well as to identify the strength and direction of the relationships between C:N:P ratios and multifunctionality.

#### Structural equation modeling

We used a structural equation modeling (SEM) approach to assess the causal relationships between factor variables, soil properties, and ecological functions. This framework allows the testing of direct and indirect pathways between the developed indices and environmental parameters. Our hypothesized causal model is based on the assumption that herbivores can affect soil structure and chemical composition (Kaštovská et al. 2024), leading to changes in the multifunctionality of soil. Thus, the main questions are whether herbivore pressure impacts soil multifunctionality and through which mediators, i.e., indirect effects. Additionally, habitat-specific characteristics are expected to influence these indices. Details on model formulations and assumptions are available in Appendix 1. Based on the results of univariate model selection and the previously formulated hypothesis, tests of directed separation (Shipley 2000) were performed. This allows the evaluation of the suitability of conceptualized pathways and the identification of missing paths. Paths without statistical support were removed from the final model, and the model fit was assessed through Fisher’s C. SEM was implemented on the *piecewise SEM* package (Lefcheck 2016) in *R software* (v4.4.1).

The workflow for the SEM approach consisted of three consecutive steps to assess the causal relationships between herbivore pressure and its direct and indirect impacts on multiple ecosystem functions. First, we assessed whether the core hypothesis supported the different pathways by examining each link’s coefficient in the fitted SEM model. Second, we examined the fit of the nested models to test the importance of each of our hypotheses in explaining our data. Furthermore, through this second step, tests of directed separation were performed to infer potential missing links, and the most parsimonious model was selected (Appendix 1).

## Results

Soil enzymatic activities varied across the herbivore pressure gradient in both seasons. Overall, soil multifunctionality was context-dependent, primarily driven by nutrient cycling and mediated by changes in physicochemical properties.

### Effects of ungulates pressure on soil enzymatic activities

Enzymatic activities showed marked seasonal variation, generally being higher in spring, with interactions between habitat and herbivore pressure. Therefore, data were analyzed separately for both seasons (Tables S2 and S3).

During fall, soil enzyme activities significantly differed across herbivore pressure and habitat (Table S2). Significant interactions between habitat and herbivore pressure were detected for dehydrogenase (DHA), acid phosphatase (AcP) activities and urease (Table S2). DHA activity increased in grasslands and forests under high herbivore pressure, but not for shrublands (Figure 2). In contrast, AcP was higher in forest control soils (Figure S3).

**Figure 2.**
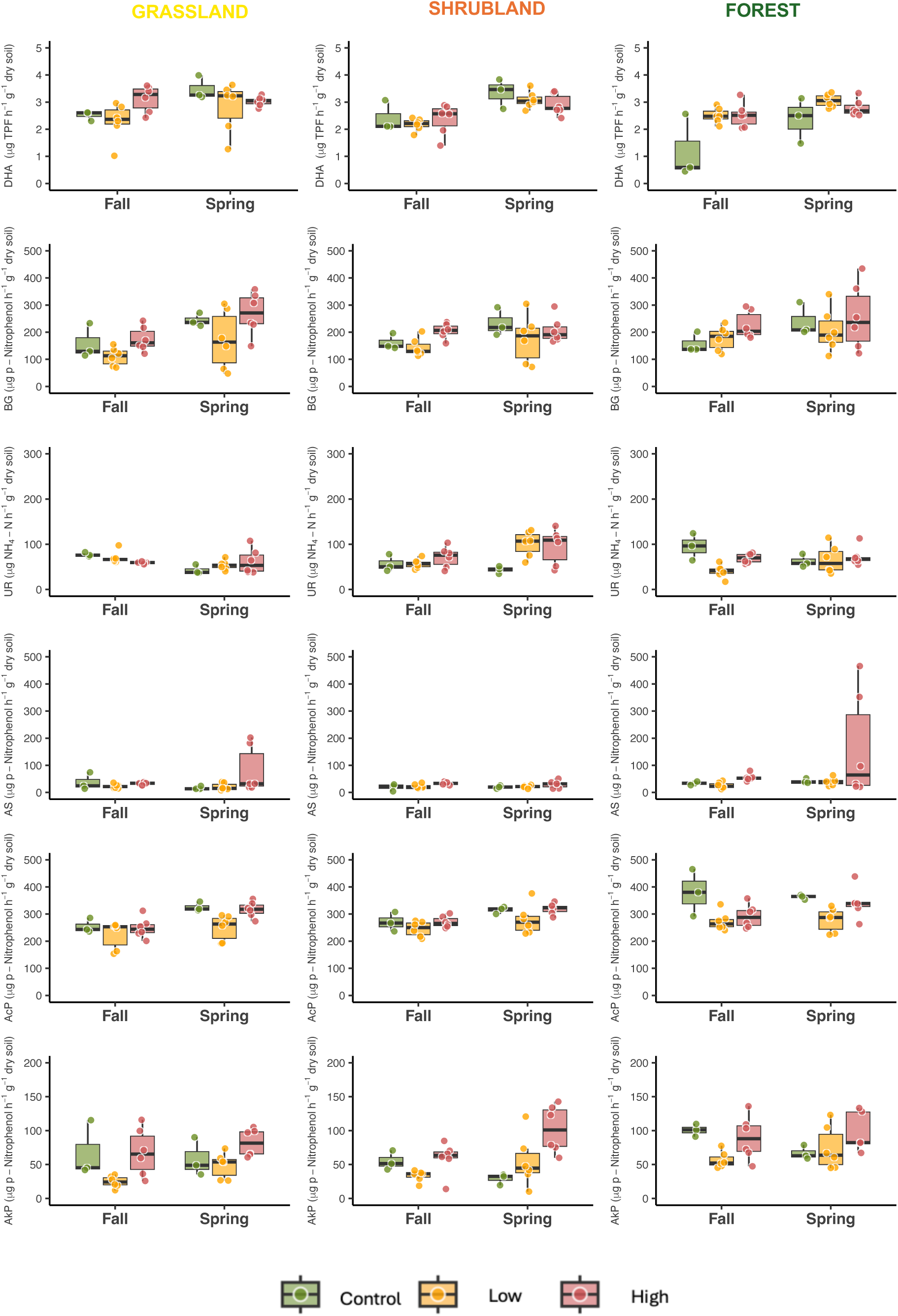
Soil enzymatic activities recorded in fall 2021 and spring 2022 for the three habitats (grasslands, shrublands and forest) within each herbivore pressure [Control (green), low (orange) and high (red) herbivore pressure].

In spring, no interactions were found between habitat and herbivore pressure (Table S2). Except for DHA, BG and UR, soil enzymatic activities significantly varied across the herbivore pressure gradient (Table S2, Figure 2). Still in spring, BG and AcP activities showed no habitat-related differences (Table S2), but both were lower at low and high herbivore pressures (Figure 2). DHA remained unaffected by habitat or herbivory pressure during spring (Table S2).

### (Multi)Functional indices are impacted by herbivore pressure

Significant correlations were found between the enzymatic activities, and soil physical and chemical properties assessed (Figure S1). Enzymatic activities assessed were also positively correlated (Figure S1), allowing their integration into a multifunctional index explaining 70% of total variance (Table S1). The multifunctional index varied across seasons, so model selection was performed separately. Low-pressure areas showed the lowest multifunctional scores across all habitats in both seasons (Figure 3). Generally, high-pressure areas showed higher multifunctionality scores than the other treatments, except during fall, in grasslands and forests (Table S2, Figure 3). Herbivore pressure and habitat explained the variation of multifunctionality in the fall. For spring, the best model only considered herbivore pressure (table S3), highlighting a lower context-specificity of habitat for this season (Figure 3). The fertility index explained 78% of the total variance (Table S1) and varied i) across habitats during spring and ii) in response to the interaction between herbivory pressure and habitats in fall (Table S2). Forests showed higher fertility values than other habitats in both seasons (Figure 3).

**Figure 3.**
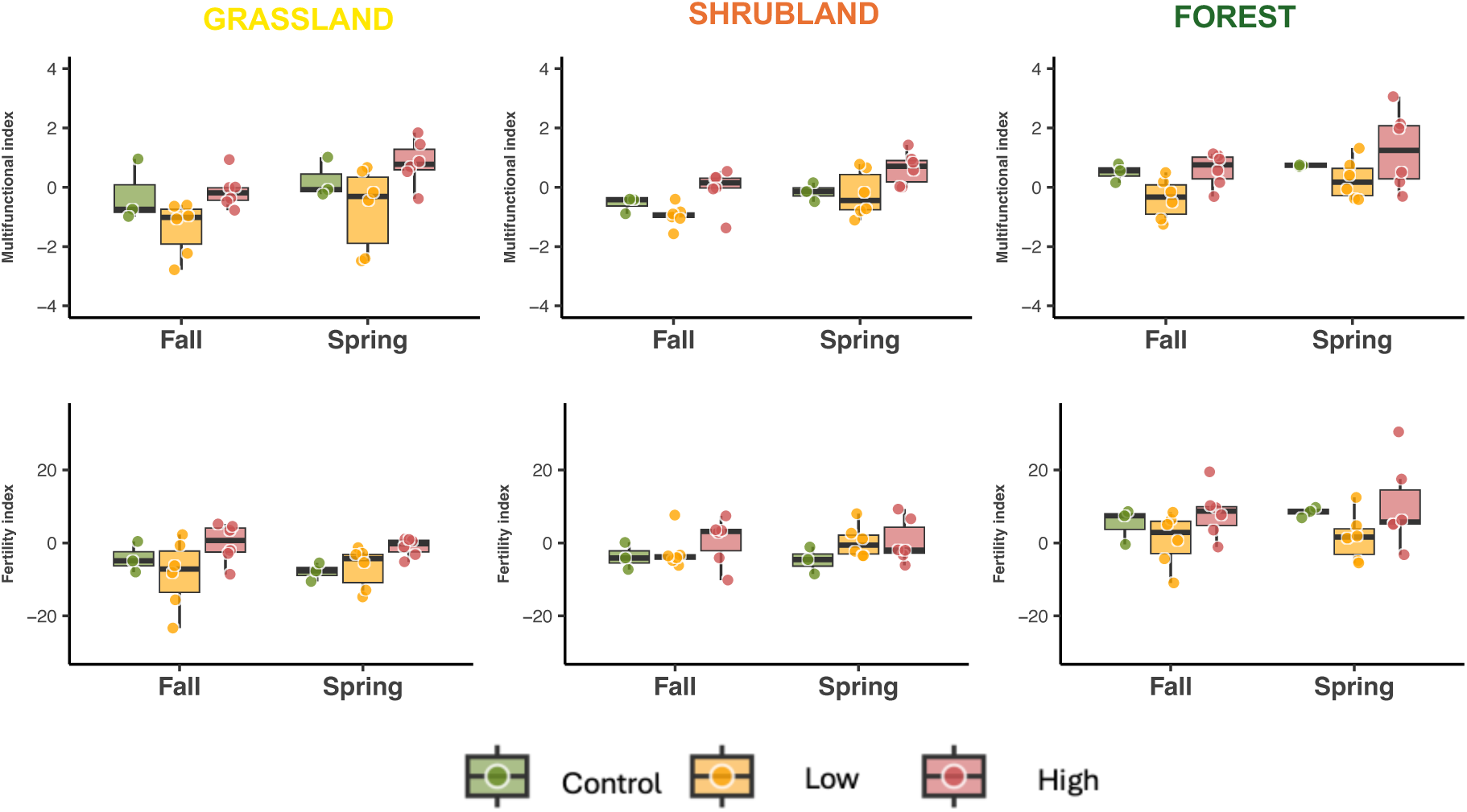
Indices for multifunctional, and fertility across each herbivore pressure [Control (no pressure), low and high pressure)], within each habitat (grasslands, shrublands, forest) during fall (2021) and spring (2022).

Indices for carbon (C), nitrogen (N), and phosphorus (P) cycling explained 76%, 64%, and 68% of the total variance (Table S1). Habitat alone best explained C cycling, whereas habitats and herbivore pressure influenced both N and P cycling (Table S3, Figure S2). The physicochemical (PCh) index explained 69% of the variance, and was driven by habitats and ungulate pressure in spring and their interaction in fall (Table S3). All functional indices correlated with the multifunctionality index, with minor distinctions in herbivore pressure treatments (Figure 4). The relationships between the indices assessed were particularly strong in forest habitats (Figure S3).

**Figure 4.**
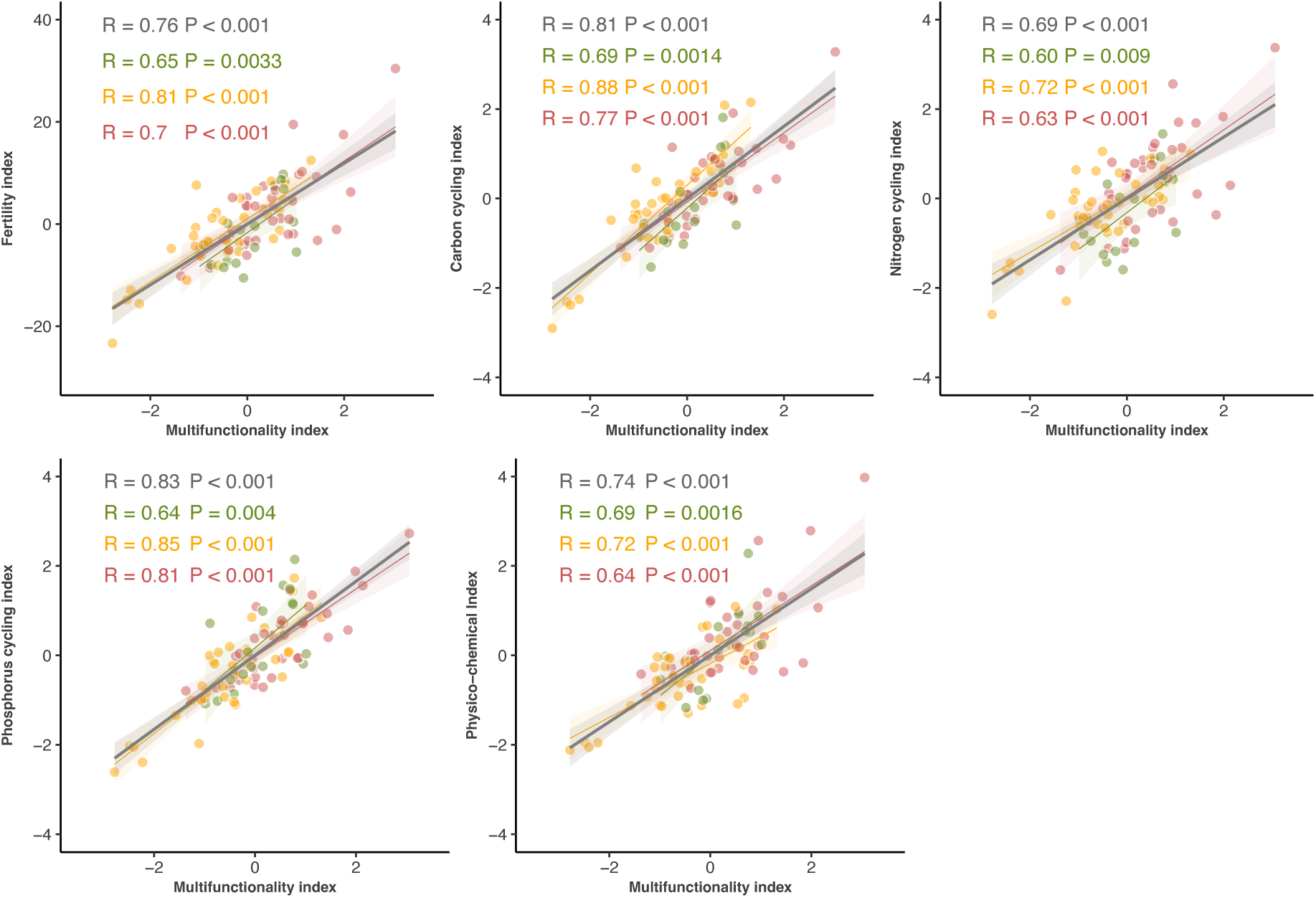
Correlations between the multifunctionality index, nutrient cycles (carbon, nitrogen, and phosphorous) and physical and chemical properties, and soil fertility indices. Points with diJerent colors correspond to diJerent herbivory pressure [Control (green), low (yellow) and high (red) herbivore pressure].

### Soil enzymatic stoichiometry determines soil multifunctionality

Soil enzyme activity stoichiometry showed mixed responses, being highly impacted by herbivore pressure (Figure 5), with a strong dependency on habitat (Figure S4).

**Figure 5.**
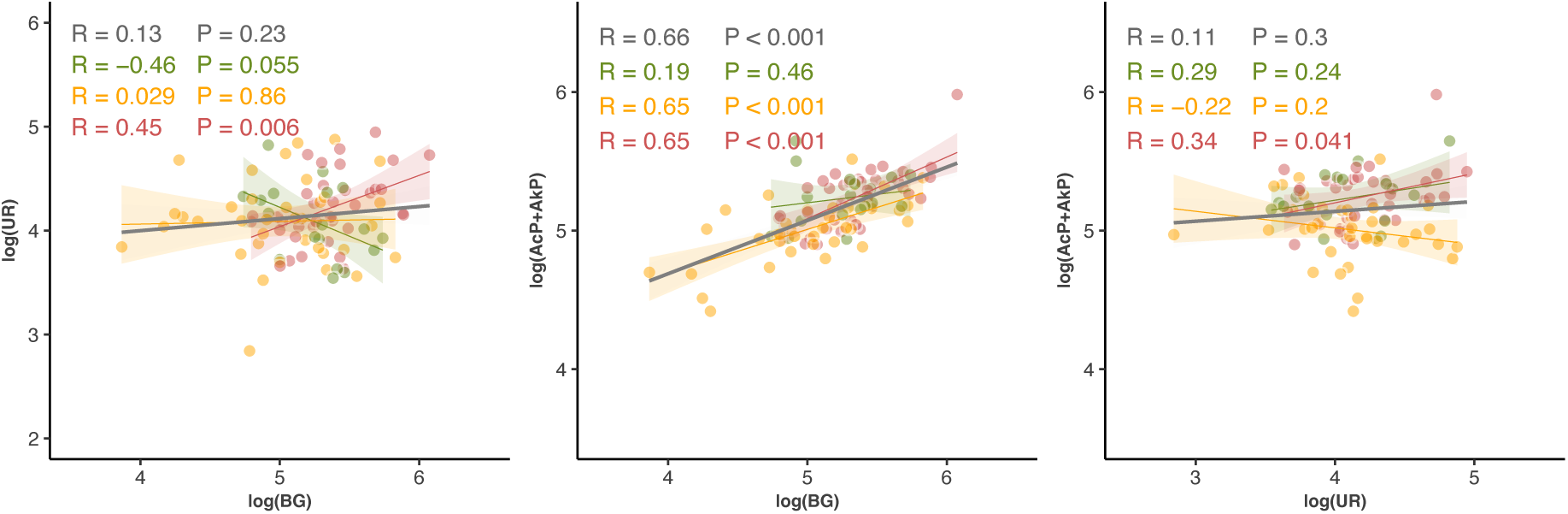
Correlation between the main enzyme activities (log transformed) responsible for each nutrient cycle (carbon – BG, nitrogen – UR and phosphorous – AcP + AkP). Points with diJerent colors correspond to diJerent herbivory pressure [Control (green), low (yellow) and high (red) herbivore pressure].

Strong positive correlations were observed between C- and P-acquiring enzymatic activities, particularly in areas subjected to low and high herbivory pressure (Figure 5). Although not evident in shrublands, these strong relationships were also reflected in grasslands and forests (Figure S4).

The C:N and N:P enzyme activity relationships proved weaker, in part due to contrasting responses between different habitats or ungulate pressures as, for instance, we found correlations of similar magnitude but opposite direction for control and high-pressure areas when regressing C against N-acquiring enzymes. A similar pattern was found for N:P relationships, with low-pressure areas denoting a contrasting correlation direction compared to the remaining.

While total N and C were tightly correlated across herbivory pressures and habitats, total nutrient stoichiometry involving P showed some context-dependent variation. Specifically, correlations between total C and P loads were moderate across herbivory pressures (Figure 6) and habitats (Figure S5). A closer insight, though, reveals that shrublands and high herbivore pressure areas contribute disproportionately to decreasing the strength of the correlations, as no significant correlations were found in these settings.

**Figure 6.**
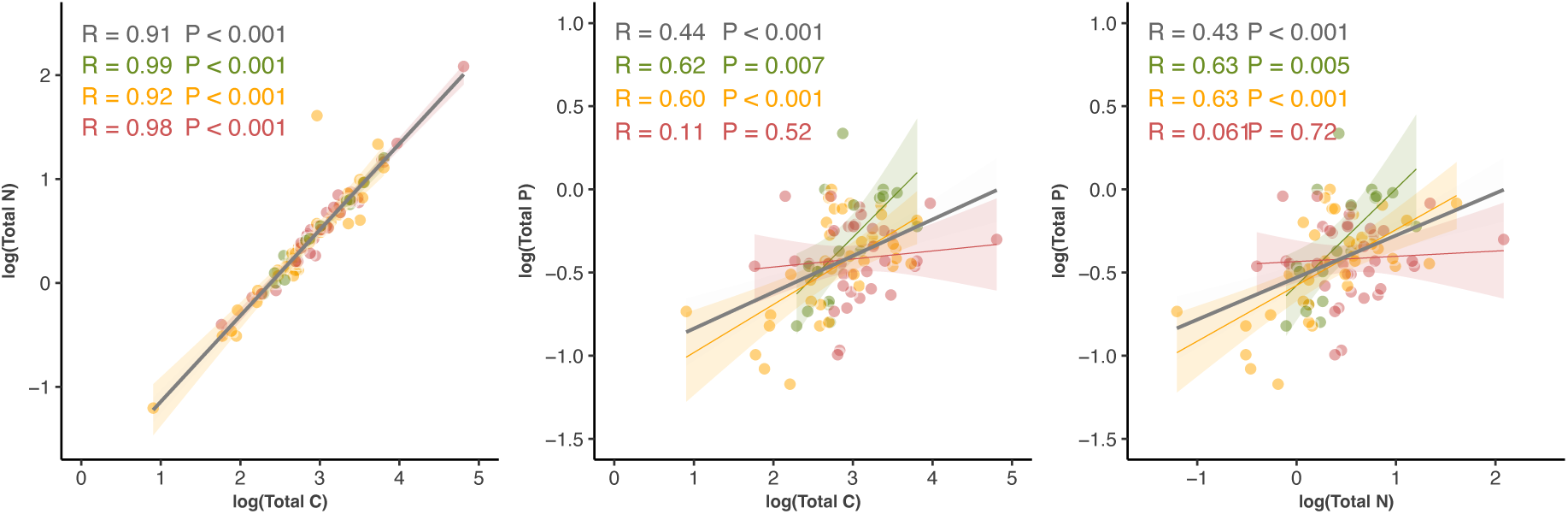
Correlation between the total carbon, nitrogen and phosphorous nutrient concentration (log transformed). Points with diJerent colors correspond to diJerent herbivory pressure [Control (green), low (yellow) and high (red) herbivore pressure].

The C:P soil enzymatic ratios showed the highest and most consistent positive correlations to the soil fertility index across herbivore pressure (Figure 7) and habitats (Figure S6). A deeper insight into the C:N and N:P enzyme activity ratios revealed mixed responses to soil fertility, which decreased the overall correlation strength. The C:N enzymatic ratio was disproportionately important to fertility at low ratio values, eventually decreasing its relevance as the ratios increased. This contrasts with the C:P enzymatic ratio, which held positive relationships over the entire range of ratio values. It is worth mentioning that the soil enzyme activity ratios were tested against the fertility index because their data are used to produce the multifunctionality index, so using the multifunctionality would inflate the correlations.

**Figure 7.**
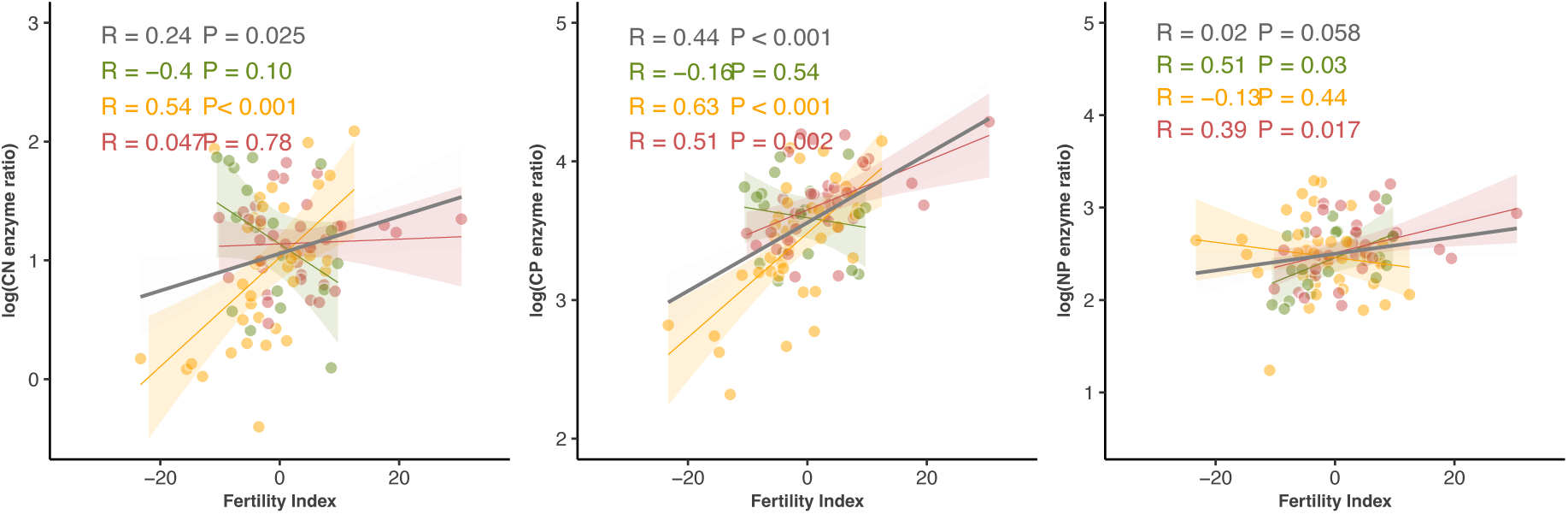
Correlation between carbon:nitrogen (CN), carbon:phosphorous (CP) and nitrogen:phosphorous (NP) enzyme ratios (log transformed) and the fertility index. Points with diJerent colors correspond to diJerent herbivory pressure [Control (green), low (yellow) and high (red) herbivore pressure].

CN, CP, and NP ratios showed strong positive relationships with soil multifunctionality particularly in forest (figure S7) and herbivore pressure (Figure 8).

**Figure 8.**
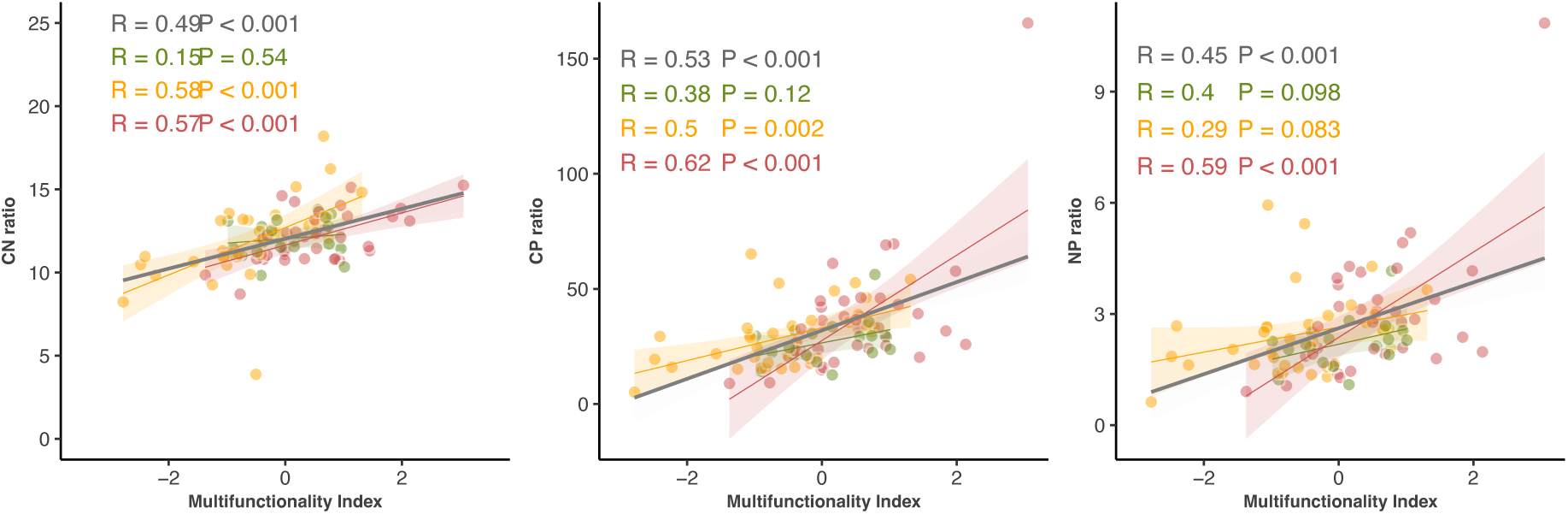
Correlation between carbon:nitrogen (CN), carbon:phosphorous (CP) and nitrogen:phosphorous (NP) enzyme ratios (log transformed) and the multifunctionality index. Points with diJerent colors correspond to diJerent herbivory pressure [Control (green), low (yellow) and high (red) herbivore pressure].

### Mechanistic pathway linking herbivory to soil processes

Structural equation models (SEM) showed that herbivore effects on soil multifunctionality were indirect, mediated by the soil properties and nutrient cycles (Figure 9). In spring, both habitat and herbivore pressure influenced soil properties (PCh index), which determined nutrient cycling, through mediated effects for C, N and P. Direct links to soil multifunctionality occurred via C and P cycling processes, aligning with the stoichiometric analysis. N cycling, did not reveal a clear link to soil multifunctionality, being its effect mediated by either PCh or C processes. The model is consistent with the strong C and N coupling, and highlights P cycling as a key of multifunctionality.

**Figure 9.**
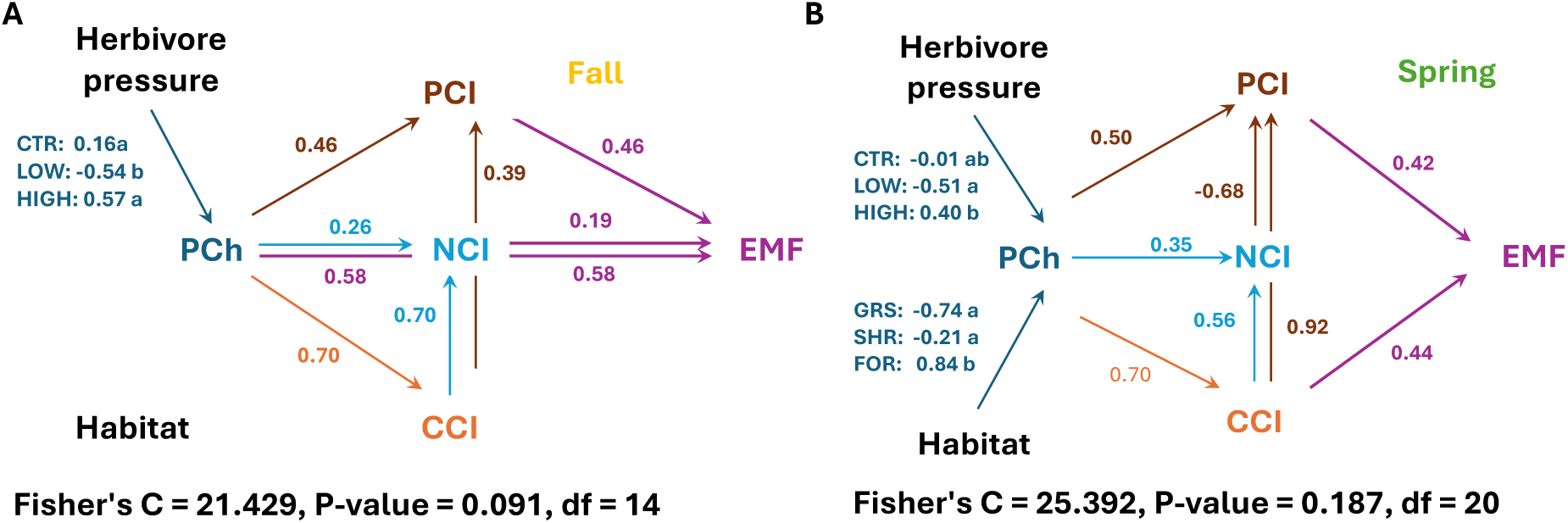
Structural equation model describing the direct and indirect eCects of ungulate herbivore pressure on multifunctionality in two seasons: Fall (A) and Spring (B). The impact of ungulates (CTR – absence, LOW – low ungulate pressure, HIGH – high ungulate pressure) is mediated by physical and chemical properties and its impacts upon carbon, nitrogen and phosphorous, described by composite indices (CCI – carbon cycle index, NCI – nitrogen cycle index, PCI – phosphorous cycle index). During spring (B), habitat characteristics (GRS – grasslands, SHR – shrublands, FOR – forests) also modulate multifunctionality, due to habitat diCerences regarding physical and chemical properties, mediating indirect eCects on nutrient cycling.

During fall, the impact of herbivores’ activity was also mediated through its influence on soil physicochemical properties and nutrient cycling. Habitat effects were not retained in the final model for fall. Soil properties (PCh) strongly affected nutrient cycles (C and P), whereas N cycling was directly affected by herbivore pressure and constituted an intermediate step to soil multifunctionality through P cycle mediation. Direct effects on multifunctionality were related to P cycling processes, but also to physical and chemical properties (Figure 9).

## Discussion

Large herbivore rewilding initiatives are increasing in Europe, underpinning the need for understanding their impact on Mediterranean landscapes. By reintroducing large herbivores, known as ecosystem engineers (Barbero-Palacios et al. 2020), these initiatives may profoundly alter soil ecosystem services (Eldridge and Soliveres 2023). Our results show that ungulates modulate soil functions linked to nutrient cycling through multiple interacting pathways, that are both strongly seasonal and habitat dependent. Beyond quantifying individual soil processes, our approach linked the effects of herbivore pressure to soil functioning, offering a practical framework for future monitoring routines.

### The impact of ungulates on soil ecological functions and its context dependency

Empirical studies have shown that ungulates influence nutrient cycling and availability, shaping plant community composition, productivity, and ecosystem functioning (Sitters and Andriuzzi 2019). Despite growing evidence suggesting their impact on soil biodiversity, particularly on microbial communities (Guo et al. 2025), few studies have examined their impacts on soil functioning.

In our study, the presence of ungulates fostered soil microbial activity, as shown by increased DHA activity (Nannipieri 2002; Wolinska and Stepniewsk 2012), but only during fall. This seasonal pattern suggests that ungulates benefit microbial communities when primary productivity is lower and resources are scarce, after the dry and hot Mediterranean summers. In addition, how ungulates use the different habitats may also explain why forests seemed to benefit from their presence while grasslands did not (Figure 2). While foraging areas such as grasslands tend to be depleted in biomass and nutrients, resting areas, such as forests, are enriched with nutrient deposition (Riesch et al. 2025).

Our results suggest that soil functional response to herbivore pressure depends on multiple interacting factors such as habitat, the intensity and space-use dynamics of the herbivores, the season and abiotic conditions, and intrinsic soil community composition. These interactions produced contrasting and context-specific enzyme activity effects. For example, during fall, how herbivore pressure impacted UR activity was modulated by habitat. While in shrublands there was a slight increase of its activity in high herbivore densities, in forests the presence of herbivores resulted in a significant decrease of its activity.

Indeed, it has been shown that UR activity is strongly impacted by vegetation, and therefore by habitat, due to changes in nitrogen distribution and availability in plant communities (Li et al. 2023).

### Soil ecological processes vary across herbivore pressure

Because herbivory affects soil processes simultaneously, integrative indices are required to evaluate soil multifunctionality, fertility and key biogeochemical functions.

The multifunctional index showed a non-linear response to herbivore pressure, with habitat and season modulation. Highest multifunctionality occurred with high herbivore pressure, regardless of season or habitat, likely driven by nutrient and labile organic matter inputs that stimulate decomposition, nutrient turnover, and fertility, as well as due to soil aeration (e.g., through rooting) (Sardans and Peñuelas 2010). In contrast, low-pressure areas showed the lowest multifunctionality, particularly during fall, when low water availability after summer and, thus, low primary productivity constrain microbial activity (Sardans and Peñuelas 2010; Hobbie and Villéger 2015). During spring, higher water availability and primary productivity enhanced multifunctionality, emphasizing the strong seasonality of Mediterranean systems. Soil multifunctionality was highly correlated with the soil fertility index, based on a large set of soil quality indicators, highlighting its potential as a practical, integrative metric for future monitoring of herbivore effects on soil functioning.

The N and C indices did not respond to the different herbivore pressures, whereas the P index was significantly affected. While N can enter the system through multiple routes, aerial and terrestrial, P derives mostly from mineral weathering (Peñuelas et al. 2013; He et al. 2020; Liu et al. 2023). Liu et al. (2023) and He et al. (2020) showed that grazing decreases the soil N and promotes soil P cycling by enhancing the activity of soil microbial communities. This finding is critical for P-limited ecosystems such as the Mediterranean (Recena et al. 2018). The C:P enzymatic ratio exhibited a strong positive relationship with soil fertility across its full range, with two key implications: (i) low ratio values indicate a greater metabolic investment in P-acquiring enzymes, but corresponded to low multifunctionality scores; (ii) high ratio values were linked to higher multifunctionality, yet followed the same linear trend, suggesting that P limitation is not entirely alleviated. In comparison, the relationship between the C:N enzymatic ratio resembled a saturating function, with increasing C:N enzymatic ratio associated with increasing soil multifunctionality up to a threshold, which remained stable. This suggests that soil multifunctionality improves below the threshold as N sufficiency increases, and lower activity of N-acquiring enzymes is required. Above this threshold, the system can no longer respond to changes in N availability, indicating other factors as more relevant (i.e., P limitation). Our results indicate that soil multifunctionality is tightly linked to nutrient cycling and fertility. Large herbivores, by (re)distributing nutrients and changing stoichiometric balances, play a central role in sustaining multifunctional soils and enhancing ecosystem resilience in Mediterranean landscapes.

### Herbivores indirectly impact multifunctionality

Our work goes beyond empirical evidence of ungulate impacts on nutrient cycling by providing a comprehensive view of their direct and indirect effects on soil functions and their context dependency. This is particularly relevant in Mediterranean landscapes, where nutrient limitations constrain ecosystem functioning. Not all nutrient dynamics were directly affected by herbivores, yet strong interactions among C, N, and P indices contributed to overall multifunctionality. Ungulate effects on EMF were primarily mediated by C and P. The deficiency of P, a limiting nutrient (Vitousek et al. 2010), is expected to severely impair multifunctionality, a relationship supported by our data. These effects were seasonal and habitat dependent. This may have arisen due to increased differences between them during spring, particularly in forests, which exhibited higher physical and chemical index scores. Additionally, phenological changes in plants, especially in forests (Richardson et al. 2010) are exacerbated during spring productivity peaks.

Impacts on C were linked to soil pH and organic matter content, through the deposition of fecal matter and trampling (Marks et al. 2024). Increased trampling activity at higher herbivore densities, diminishes soil aggregates and accelerates organic matter decomposition (Neff et al. 2005). This may lead to an increase in carbon stocks through organic matter deposition, and an increase in phosphorus mineralization and thus, nutrient availability.

The effect of ungulates on multifunctionality was indirect, via their indirect influences on nutrient cycling, especially of C and P. During fall, the impacts of herbivores on multifunctionality are mediated by their effects on physical and chemical properties, modulating and, in particular, on phosphorous cycling. In Spring, the impacts of large herbivores are mediated through P and C cycling, due to direct impacts on physical and chemical properties. Areas with high herbivore pressure showed improved soil moisture, pH, bulk density and organic matter, regardless of the season. Nonetheless, the impact of these parameters was more important during fall as it shows direct effects on multifunctionality. Our results are consistent with previous studies, as increases in soil organic carbon can occur through mechanical actions in soil (Wei et al. 2021) or by deposition of excretions (Wang et al. 2022).

Herbivores tend to select for plant species richer in nitrogen (higher quality) (Ritchie et al. 1998), which in turn shape soil physical and chemical characteristics that may enhance nutrient cycling (e.g., soil mixing and aeration) and contribute to an input of organic matter providing both carbon and nitrogen. Herbivory can change the stoichiometry of nutrients by impacting each nutrient cycle differently (He et al. 2020). It has been noted that higher herbivore pressure may produce positive outcomes by promoting a higher mixture of soil with excretion and greater rates of N fixation enhancing microbial diversity and activity and stimulating the decay rate of litter (Knops, 2002). On the contrary, another study suggests that imbalance in C:N ratios may be produced through a decrease in soil organic matter, when herbivores are absent (Wang et al. 2014). In future studies, the inclusion of nitrogen fractions may aid in understanding which stages of organic matter decomposition would be predominant in these soils. Regarding phosphorous ratios, grazing may increase leaf N:P ratios in many plant functional groups, suggesting that P could become more important for plant productivity at the community level (Yu et al. 2021). The ability of herbivores to shape C:N:P stochiometric and consequently modulate microbial community composition and enzymatic activities (Shen et al. 2019) is key for soil functioning and the provision of services. From this perspective, our work further highlights the pivotal role of large herbivores in regulating soil biogeochemistry and supporting ecosystem services. Their influence on nutrient dynamics and soil functioning underscores their importance in maintaining ecosystem multifunctionality across diverse habitats.

### Implications for landscape management

Mediterranean ecosystems are undergoing rapid global changes, including increasing aridity, which threatens the many ecosystem services they provide (Peñuelas et al. 2017). Although large herbivores are integral elements of these landscapes, their effects on the soil belowground processes remain poorly understood. With current rewilding efforts aimed at restoring trophic webs, understanding their influence on soil functioning is essential to promote biodiverse and self-sustaining Mediterranean landscapes. Despite the growing recognition of the importance of soil ecosystem services in science and policy, standardized indicators for their assessment are still lacking. Establishing baseline values for healthy soils is a major challenge largely due to the intrinsic heterogeneity and strong context-dependence of these ecosystems, across spatial and temporal scales. Our results highlight this variability emphasizing the need to assess soil ecological functions across contexts to develop comprehensive indicators with broader ecological relevance.

The ecological indices proposed here, encompassing soil ecological functions related to biogeochemical cycles, were found to be sound predictors, not only of the impacts of herbivory across a range of habitats, but also responsive to soil properties at a finer scale. This implies that, within the contexts of this landscape, these indices may be applied to other landscape use scenarios, for predicting changes in soil ecological functions.

Although more studies and empirical data are needed to fully understand how large herbivores impact the soil compartment, the indices produced in this study are promising and supportive tools bridging conservation and the provision of ecosystem services. In the context of rewilding initiatives, much has been advocate regarding the role of large herbivores in carbon sequestration and persistence (Kristensen et al. 2022). Here, we provide evidence of the role of large herbivores on the engineering of nutrient cycling as whole, not just carbon, with the ability to fully shape landscapes bottom-up.

## Author contribution statement

**Jorge F. Henriques**: Conceptualization, Methodology, formal analysis, data curation, visualization, writing – Original draft; **Rui G. Morgado**: Conceptualization, Methodology, sofware, visualization, writing – Review & editing, funding acquisition, supervision; **João Carvalho**: Conceptualization, Methodology, Visualization, writing review & editing, funding acquisition, supervision, project administration; **Rita Tinoco Torres**: Conceptualization, writing review & editing, funding acquisition, supervision,, funding acquisition, **Mariana Rossa**: Methodology, writing – Review & editing, **Joana Fernandes**: Methodology, writing – Review & editing, **Catarina Malheiro**: Methodology, writing – Review & editing, **Marija Prodana**: Methodology, writing – Review & editing, **Sara Peixoto**: Methodology, writing – Review & editing, **Eduardo Ferreira**: Methodology, writing – Review & editing, **Susana Loureiro**: Conceptualization, writing – Review & editing, **Jennifer A. Krummins**: Conceptualization, writing – Review & editing; **Ramón Perea**: Conceptualization, writing – Review & editing; **Emmanuel Serrano**: Conceptualization, writing – Review & editing.

