## supplementary figures 1-9 for "Herbivore pressure modulates soil multifunctionality in a Mediterranean landscape"

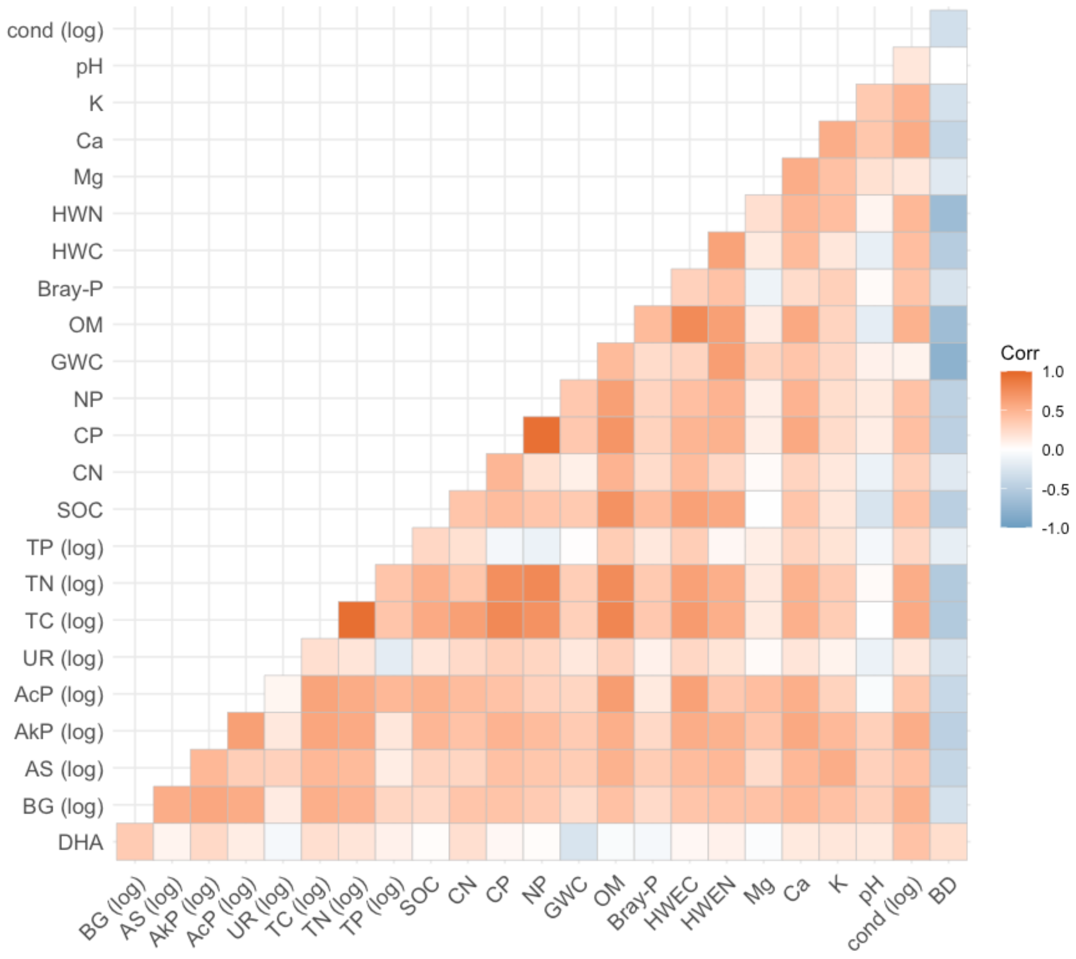


Figure S1 – Heatmap representing the correlation coefficients between enzymes activity and soil physical and chemical properties. DHA – dehydrogenase; BG - β-glucosidase; AS – arylsulfatase; AkP - alkaline phosphatase; AcP - acid phosphatase; UR – urease; TC – total carbon; TN – total nitrogen; TP – total phosphorous; TH – total hydrogen; SOC – soil organic carbon; CN – C:N ratio; CP – C:P ratio; NP – N:P ratio; GWC – gravimetric water content; OM – organic matter; Bray-P – extractable (Bray-1) phosphorous; HWC – hot water extractable carbon; HWN – hot water extractable nitrogen; Mg – magnesium; Ca – calcium; K – potassium; cond – conductivity; BD – bulk density.

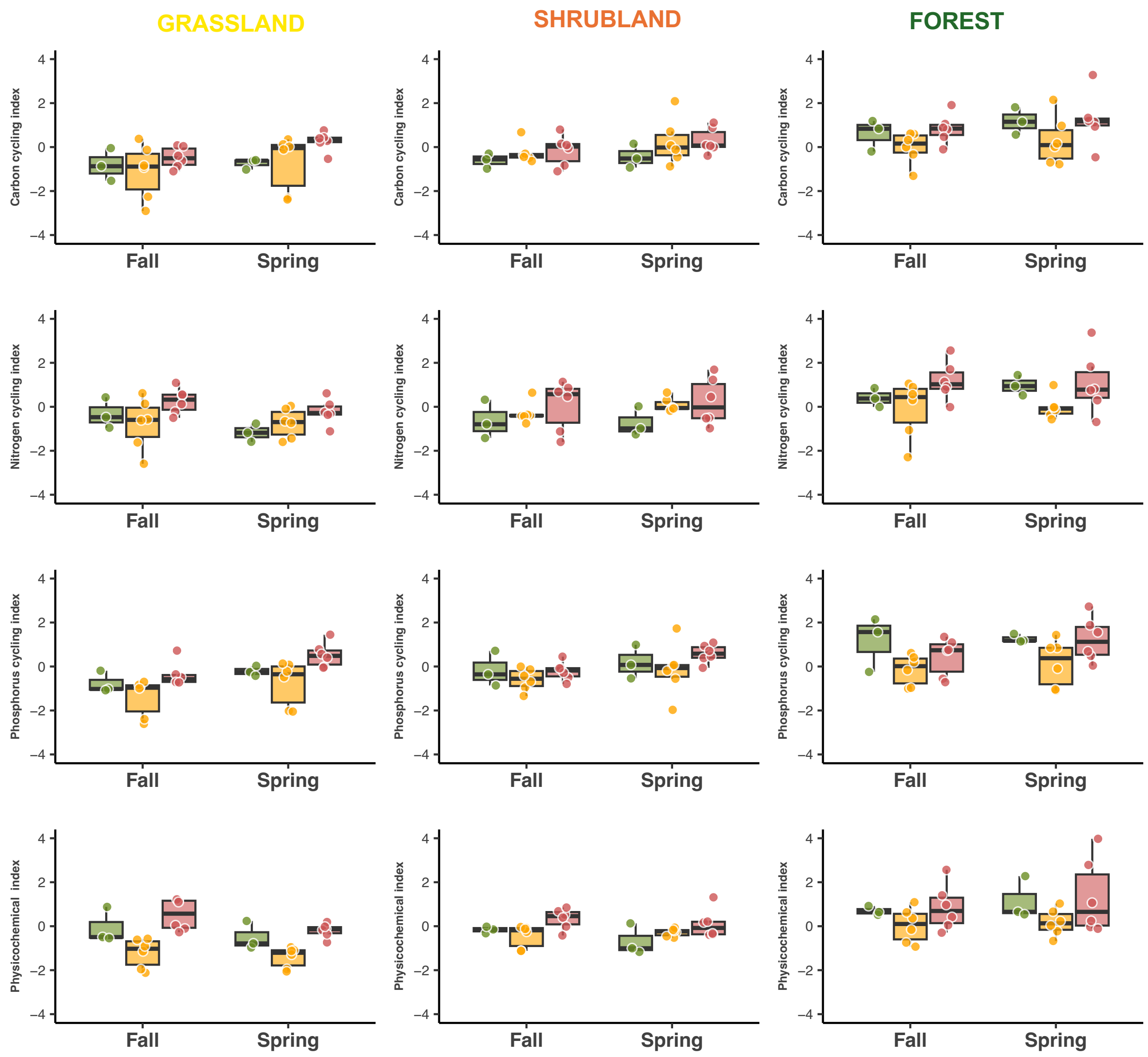


Figure S2 – Soil nitrogen cycling indices recorded in fall 2021 and spring 2022 for the three habitats (grasslands, shrublands and forest) within each herbivore pressure [Control (green), low (orange) and high (red) herbivore pressure].

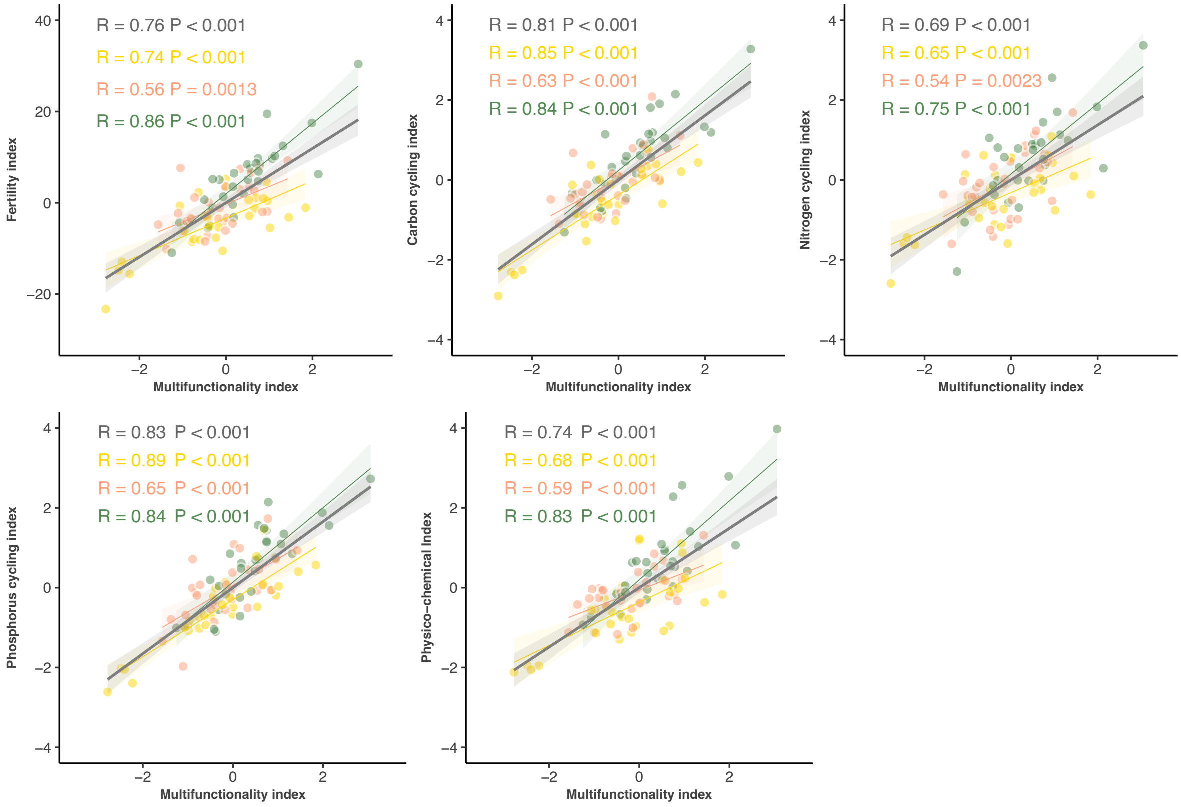


Figure S3 – Correlations between multifunctionality index with soil fertility indices and each nutrient cycle index (carbon, nitrogen, and phosphorous). Points with different colors correspond to different habitats [grasslands (yellow), shrublands (orange) and forests (green) herbivore pressure].

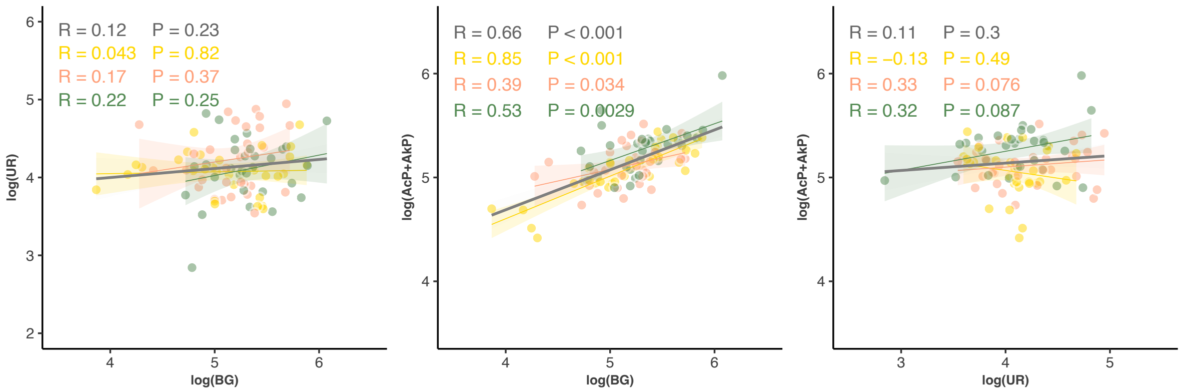


Figure S4 – Correlation between the main enzyme activities (log transformed) responsible for each nutrient cycle (carbon – BG, nitrogen – UR and phosphorous – AcP+ AkP). Points with different colors correspond to different habitats [grasslands (yellow), shrublands (orange) and forests (green) herbivore pressure].

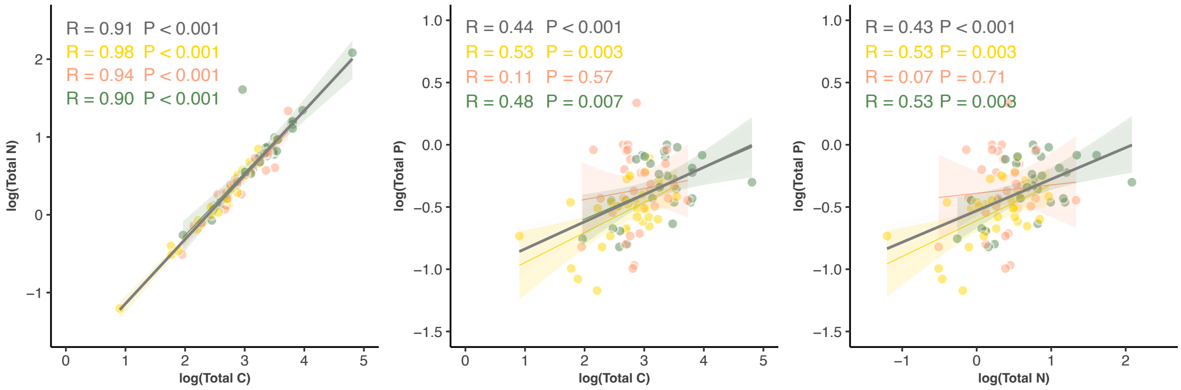


Figure 5S – Correlations between the total carbon, nitrogen and phosphorous nutrient concentration (log transformed). Points with different colors correspond to different habitats [grasslands (yellow), shrublands (orange) and forests (green) herbivore pressure].

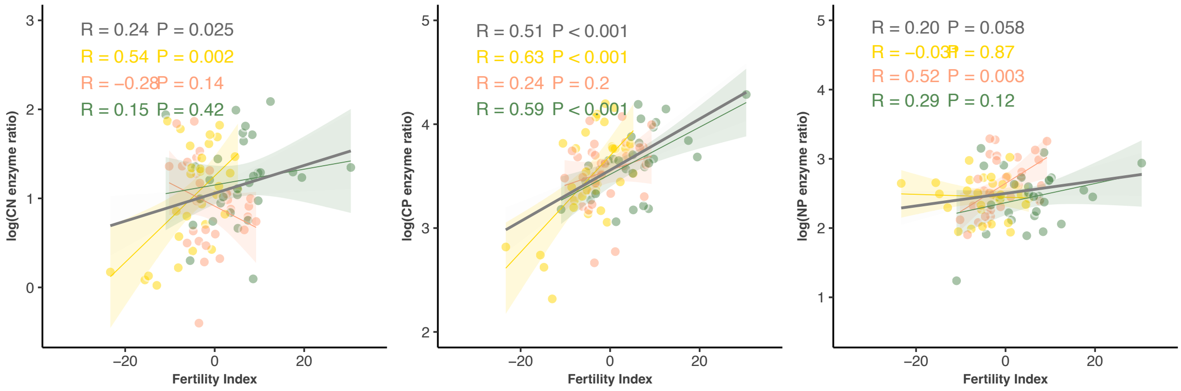


Figure S6 – Correlations between carbon:nitrogen (CN), carbon:phosphorous (CP) and nitrogen:phosphorous (NP) enzyme ratios (log transformed) and the fertility index.
Points with different colors correspond to different habitats [grasslands (yellow), shrublands (orange) and forests (green) herbivore pressure].

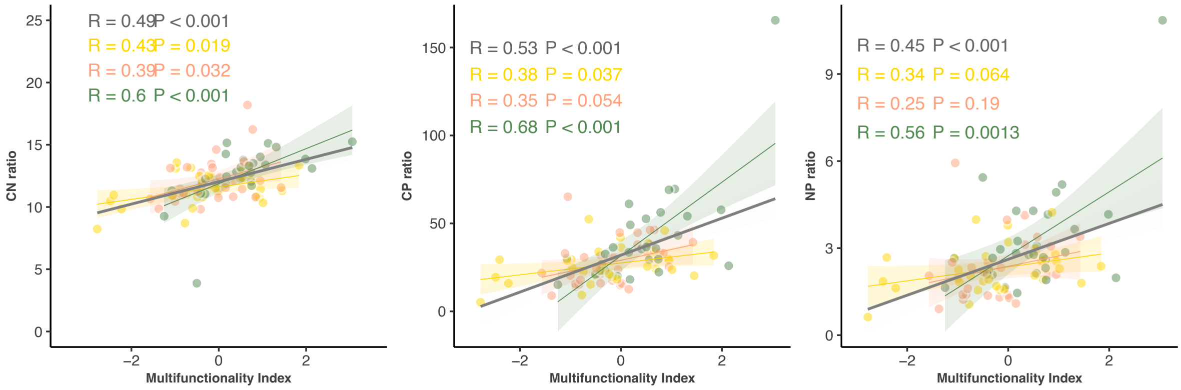


Figure S7 – Correlations between carbon: nitrogen (C:N), carbon: phosphorous (C:P) and nitrogen: phosphorous (N:P) enzyme ratios (log transformed) and the multifunctionality index. Points with different colors correspond to different habitats [grasslands (yellow), shrublands (orange) and forests (green) herbivore pressure].

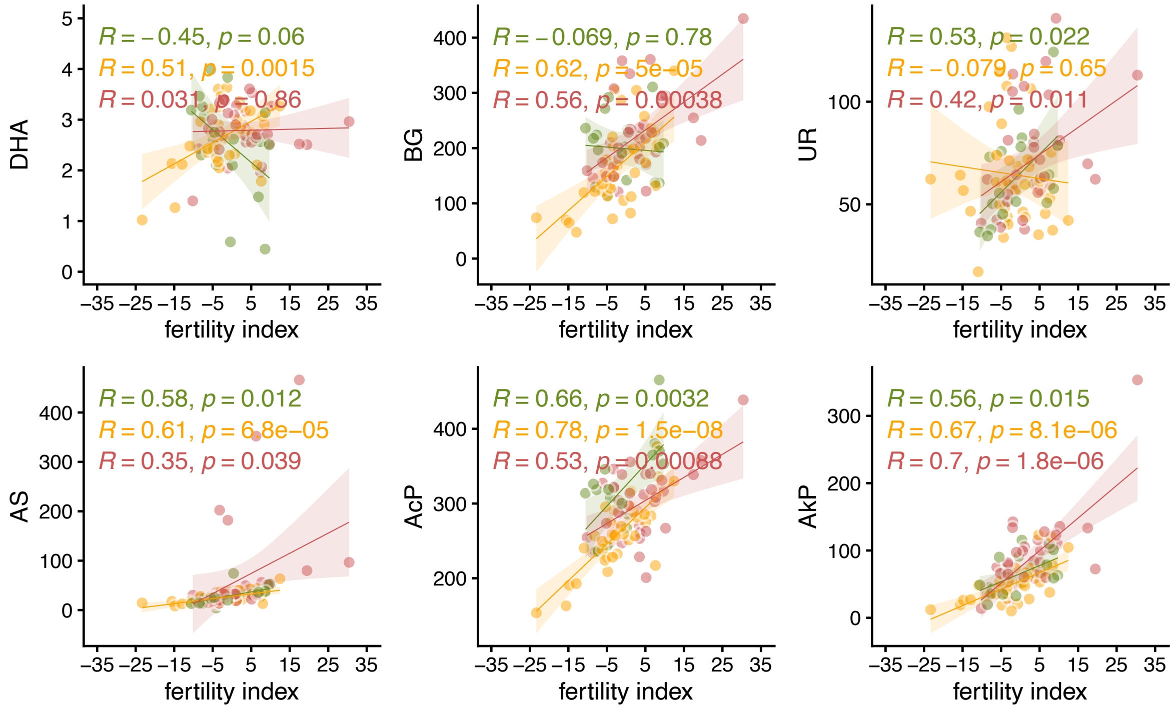


Figure S8 – Correlations between each enzyme activity and the fertility index.
DHA – dehydrogenase; BG - β-glucosidase; AS – arylsulfatase; AkP - alkaline phosphatase; AcP - acid phosphatase. Points and lines with different colors correspond to different herbivore pressure [Control (green), low (yellow) and high (red) herbivore pressure].

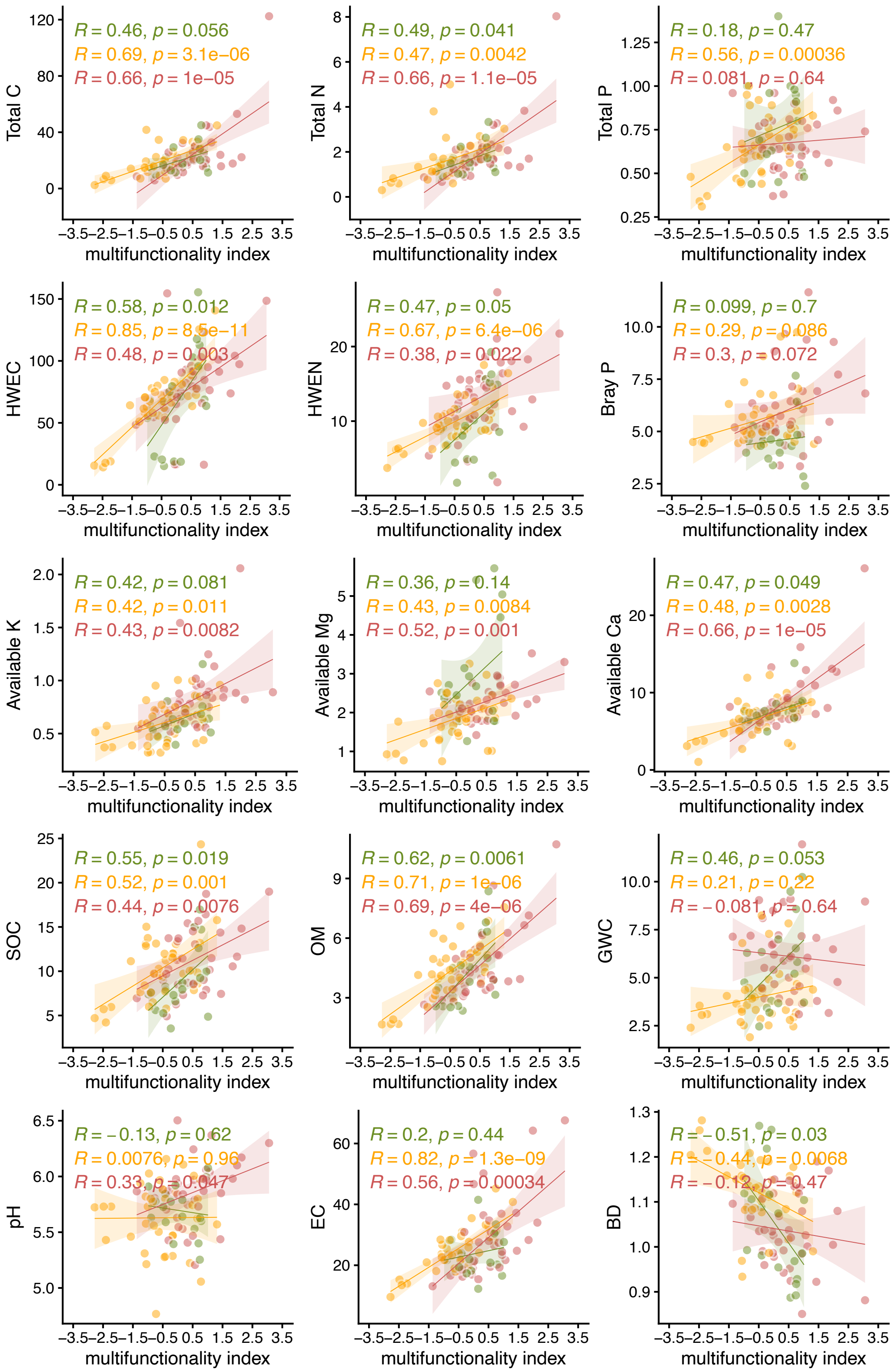


Figure S9 - Correlations between each enzyme activity and the fertility index. DHA – dehydrogenase; BG - β-glucosidase; AS – arylsulfatase; AkP - alkaline phosphatase; AcP - acid phosphatase. Points and lines with different colors correspond to different herbivore pressure [Control (green), low (yellow) and high (red) herbivore pressure].
