## Supplementary tables 1-3 for "Herbivore pressure modulates soil multifunctionality in a Mediterranean landscape"

Table S1 – Principal component analysis (PCA) loadings and scores obtained for each one of the multifunctionality and nutrient cycling indices.

| | Index | | | | | |
|---|---|---|---|---|---|---|
| Variable | Multifunctionality | Nitrogen | Carbon | Phosphorous | Physico-chemical | Fertility |
| DHA | 0.36 | | | | | |
| b-glucosidase (ln) | 0.85 | | 0.65 | | | |
| Aryl-sulf (ln) | 0.71 | | | | | |
| Alkaline phos (ln) | 0.71 | | | 0.79 | | |
| Acid phosphatase | 0.76 | | | 0.88 | | |
| Urease (ln) | 0.28 | 0.47 | | | | |
| Nitrogen (ln) | | 0.84 | | | | 0.88 |
| Carbon (ln) | | | 0.88 | | | 0.91 |
| Phosphorous | | | | 0.62 | | 0.29 |
| SOC | | | 0.78 | | | 0.70 |
| OM | | | | | 0.73 | 0.90 |
| pH | | | | | 0.28 | 0.03 |
| Conductivity (ln) | | | | | 0.67 | 0.65 |
| GWC | | | | | -0.76 | 0.54 |
| BD | | | | | 0.64 | -0.72 |
| Bray-P | | | | 0.43 | | 0.49 |
| HWEC | | | 0.86 | | | 0.75 |
| HWEN | | 0.84 | | | | 0.77 |
| CN | | | | | | 0.53 |
| CP (ln) | | | | | | 0.81 |
| NP (ln) | | | | | | 0.74 |
| Mg | | | | | 0.54 | 0.26 |
| Ca | | | | | 0.84 | 0.67 |
| K | | | | | 0.68 | 0.47 |
| Cumulative (%) | 0.70 | 0.64 | 0.76 | 0.68 | 0.69 | 0.78 |

Dehydrogenase (DHA), β-glucosidase (BG), aryl sulfatase (AS), acid phosphatase (AcP), alkaline phosphatase (AkP), urease (UR), SOC – soil organic carbon, CN – Carbon/nitrogen ratio, OM – organic matter, GWC – gravimetric water content, BD – Bulk density, CP – carbon/phosphorous ratio, NP – nitrogen/phosphorous ratio.

Table S2 – Model selection for the impact of herbivore pressure and habitat on the soil enzymes assessed for two seasons (fall 2021 and spring 2022: dehydrogenase (DHA), β-glucosidase (BG).aryl sulfatase (AS), acid phosphatase (AcP), alkaline phosphatase (AkP) and urease (UR), Bold denotes statistical significant differences, hab – habitat, her - herbivore pressure.

| FALL | DHA | | BG | | AS (ln) | | AcP | | AkP | | UR (ln) | |
|---|---|---|---|---|---|---|---|---|---|---|---|---|
| | AIC | ΔAIC | AIC | ΔAIC | AIC | ΔAIC | AIC | ΔAIC | AIC | ΔAIC | AIC | ΔAIC |
| fixed effects | | | | | | | | | | | | |
| null | 90.42829 | 0 | 475.2093 | 0 | 61.28666 | 0 | 483.3046 | 0 | 78.5122 | 0 | 22.05982 | 0 |
| hab | 91.03985 | 0.61156 | 475.255 | 0.0457 | 61.64843 | 0.36177 | 476.7279 | -6.5767 | 74.31219 | -4.20001 | 25.17381 | 3.11399 |
| dens | 89.2298 | -1.19849 | 469.8952 | -5.3141 | **58.69137** | **-2.59529** | 481.7572 | -1.5474 | 70.45237 | -8.05983 | 20.66411 | -1.39571 |
| hab + her | 89.16174 | -1.26655 | **467.6036** | **-7.6057** | 57.86543 | -3.42123 | **472.2567** | **-11.0479** | **60.27851** | **-18.23369** | 23.58133 | 1.52151 |
| hab + her + hab:her | **83.936** | **-6.49229** | 472.6394 | -2.5699 | 61.31848 | 0.03182 | 473.3361 | -9.9685 | 64.72409 | -13.78811 | **15.4816** | **-6.57822** |

| SPRING | DHA | | BG | | AS (ln) | | AcP | | AkP | | UR (ln) | |
|---|---|---|---|---|---|---|---|---|---|---|---|---|
| | AIC | ΔAIC | AIC | ΔAIC | AIC | ΔAIC | AIC | ΔAIC | AIC | ΔAIC | AIC | ΔAIC |
| fixed effects | | | | | | | | | | | | |
| null | 54.5742 | 0 | 531.5228 | 0 | 107.9006 | 0 | 475.5986 | 0 | 82.07867 | 0 | 37.19005 | 0 |
| hab | 55.73 | 1.1558 | 534.359 | 2.8362 | 105.3831 | -2.5175 | 477.4239 | 1.8253 | 82.09533 | 0.01666 | 36.74521 | -0.44484 |
| dens | 58.1565 | 3.5823 | 530.4861 | -1.0367 | 106.6085 | -1.2921 | **464.2999** | **-11.2987** | 73.53149 | -8.54718 | 35.77001 | -1.42004 |
| hab + her | 59.26943 | 4.69523 | 533.0402 | 1.5174 | **102.5597** | **-5.3409** | 464.6057 | -10.9929 | **71.21222** | **-10.86645** | 34.25068 | -2.93937 |
| hab + her + hab:her | 60.2698 | 5.6956 | 539.6707 | 8.1479 | 108.8323 | 0.9317 | 470.6485 | -4.9501 | 75.51928 | -6.55939 | 36.35653 | -0.83352 |

Table S3 - Model selection for the impact of herbivore density and habitat on the soil multifunctional indices determined for two seasons (fall 2021 and spring 2022): Multifunctional (enzymes), nitrogen (N), carbon (C), phosphorous (P), Physical and chemical (PCh) and fertility indexes. hab – habitat, her - herbivore pressure.

| FALL | Multifunctional | | Nitrogen | | Carbon | | Phosphorous | | PCh | | Fertility | |
|---|---|---|---|---|---|---|---|---|---|---|---|---|
| | AIC | ΔAIC | AIC | ΔAIC | AIC | ΔAIC | AIC | ΔAIC | AIC | ΔAIC | AIC | ΔAIC |
| fixed effects | | | | | | | | | | | | |
| null | 107.4938 | 0 | 123.5103 | 0 | 111.4783 | 0 | 113.7398 | 0 | 115.1445 | 0 | 307.1647 | 0 |
| hab | 104.8747 | -2.6191 | 123.0604 | -0.4499 | **102.0159** | **-9.4624** | 105.6462 | -8.0936 | 113.9972 | -1.1473 | 301.694 | -5.4707 |
| her | 100.2929 | -7.2009 | 127.5644 | 4.0541 | 113.4035 | 1.9252 | 113.1282 | -0.6116 | 106.9491 | -8.1954 | 306.8395 | -0.3252 |
| hab + her | **93.35112** | **-14.14268** | **120.8593** | **-2.651** | 102.4799 | -8.9984 | **101.9952** | **-11.7446** | **102.3609** | **-12.7836** | **299.3327** | **-7.832** |
| hab + her+ hab: her | 99.77647 | -7.71733 | 127.2324 | 3.7221 | 108.1836 | -3.2947 | 106.5595 | -7.1803 | 106.0772 | -9.0673 | 305.7684 | -1.3963 |

| SPRING | Multifunctional | | Nitrogen | | Carbon | | Phosphorous | | PCh | | Fertility | |
|---|---|---|---|---|---|---|---|---|---|---|---|---|
| | AIC | ΔAIC | AIC | ΔAIC | AIC | ΔAIC | AIC | ΔAIC | AIC | ΔAIC | AIC | ΔAIC |
| fixed effects | | | | | | | | | | | | |
| null | 128.5725 | 0 | 129.6461 | 0 | 136.1503 | 0 | 125.045 | 0 | 136.8099 | 0 | 318.1408 | 0 |
| hab | 129.3199 | 0.369 | 120.7089 | -8.9372 | **130.2141** | **-5.9362** | 122.7792 | -2.2658 | 124.3393 | -12.4706 | **305.5139** | **-12.6269** |
| her | **123.2937** | **-4.182** | 129.8234 | 0.1773 | 136.7526 | 0.6023 | 121.5008 | -3.5442 | 136.0206 | -0.7893 | 319.0319 | 0.8911 |
| hab + her | 122.6208 | -5.22 | **118.6083** | **-11.0378** | 129.3634 | -6.7869 | **117.0746** | **-7.9704** | **118.0659** | **-18.744** | 303.7454 | -14.3954 |
| hab + her+ hab: her | 129.1884 | 1.208 | 119.5849 | -10.0612 | 132.9375 | -3.2128 | 123.4325 | -1.6125 | 121.2364 | -15.5735 | 306.1406 | -12.0002 |
